# PyBootNet: A Python Package for Bootstrapping and Network Construction

**DOI:** 10.1101/2024.08.08.607205

**Authors:** Shayan R. Akhavan, Scott T. Kelley

**Affiliations:** Bioinformatics and Medical Informatics Program, San Diego State University, San Diego, CA, USA; Department of Biology, San Diego State University, San Diego, CA, USA

## Abstract

PyBootNet is a user-friendly Python package that integrates bootstrapping analysis and correlation network construction. The package offers functions for generating bootstrapped network metrics, statistically comparing network metrics among datasets, and visualizing bootstrapped networks. PyBootNet is designed to be accessible and efficient, with minimal dependencies and straightforward input requirements. To demonstrate its functionality, we applied PyBootNet functions to compare networks within two disparate microbial community datasets: a mouse gut microbiome study and a microbiome study of a built environment. The PyBootNet functions applied include data preprocessing, bootstrapping, correlation matrix calculation, network statistics computation, and network visualization. In both datasets, we show that PyBootNet can generate robust bootstrapped network metrics and identify significant differences in one or more network metrics between pairs of networks. We also show that PyBootNet can create bootstrapped network graphs and identify clusters of nodes that are highly interconnected. We also confirmed its computational efficiency and scalability, which allows it to handle large and complex datasets. PyBootNet provides a powerful and extendible Python bioinformatics solution for bootstrapping analysis and network construction that can be applied to microbial, gene, metabolite and other biological data appropriate for network correlation comparison and analysis.

## INTRODUCTION

Network analysis has emerged as a powerful statistical tool for investigating and interpreting complex interactions within microbial communities. The construction of networks employs mathematical algorithms to model and visualize the relationships between different microbial species, providing valuable insights into ecosystem structure, dynamics, and functional roles [1]. By applying network analysis techniques to microbial community data, researchers can uncover interaction patterns among microorganisms [2]. Data science and the development of bioinformatics have revolutionized many fields, including the study of microbial communities. Understanding microbial community structure is crucial for advancing clinical research and gaining insights into the complex interactions within observed communities [3].

A network consists of circular nodes connected by lines, also known as edges. In microbial communities, nodes typically represent different microbial taxa, while edges indicate the strength and directionality of interactions between nodes. A common metric for establishing edge strength is the statistical correlation between nodes (e.g., Spearman or Pearson) which can be visually represented via the length, color, and/or width of edge lines. The number of edges connected to a node, known as its degree, provides valuable insights into the potential interactions between a specific microbe and others within the community, helping researchers understand the complex dynamics of microbial ecosystems [5]. For example, in the human oral microbial community positive correlations between *Streptococcus* and *Veillonella* suggested a potential metabolic relationship where *Streptococcus* produces lactate that *Veillonella* consumes [6]. In contrast, negative correlations could indicate competition for similar resources to survive, such as *Bacteroides* and *Bacilli* in the oral cavity [7]. Alternatively, strong correlations also indicate that organisms are subject to the same underlying environmental conditions, i.e., identical growth conditions. Other metrics such as betweenness centrality and transitivity describe the architecture of networks and can be used to infer potential biological characteristics. For example, nodes with high betweenness centrality, which calculate the number of shortest paths passing through nodes, serve as valuable information for prioritizing nodes as potential therapeutic targets when identifying central transcription factors and post-translational proteins[8]. Additionally, high transitivity within the network indicates the existence of closely linked node clusters and the division of the network into separate subcomponents that provide insight into the stability of the community structures existing between species [9].

Two aspects of networks that are often ignored or difficult to assess are (1) the statistical robustness of the network, and (2) if networks from two different biological systems or experimental conditions are statistically different from one another. Indeed, many papers describe networks and network metrics without any statistical tests. One method that holds special promise in the statistical evaluation of networks is the bootstrap [24]. Bootstrapping is a resampling technique that estimates the sampling distribution of a statistic and constructs confidence intervals. This method is particularly useful when the underlying distribution of the test statistic is unknown, when the dataset is self-contained (all the sample data is used to generate the statistic) and when dealing with population parameters other than the mean [3]. Such a method appears to be ideal for estimating confidence intervals for networks. With networks, all the samples are used to create the networks, and network metrics such as betweenness centrality and transitivity have unknown distributions. Bootstrap sampling distributions provide valuable information about the variability and uncertainty associated with a given statistic. This information is used to construct confidence intervals, which offer a range of plausible values for the true population parameters based on the observed sample data. By employing bootstrapping techniques, researchers can make more robust inferences even when faced with limited sample sizes or complex underlying distributions [10].

Several packages and modules, such as BioNetComp [11] in Python and BootNet [12] in R, have been developed to apply bootstrapping to the statistical analysis of correlation networks. While useful, these modules present certain accessibility issues. For example, BioNetComp requires input data with a reference database to construct accurate network metrics. This proves challenging or impossible when dealing with counts of uncultured microbial taxa from an environmental community, or untargeted metabolomic data. Indeed, BioNetComp was designed specifically for interactomes from differentially expressed genes and is not generalizable for other types of datasets. The R package BootNet performs bootstrap analysis with any given properly formatted feature table. However, its numerous dependencies have led to version control issues, requiring end-users to seek assistance running simple metrics. Moreover, BootNet focuses on examining the statistical robustness within networks and does not have features for comparing networks from different conditions or environments.

Here, we describe the development of PyBootNet, a flexible network bootstrapping package in Python that provides simple and intuitive functions for generating numerous bootstrapped network metrics, statistically comparing network metrics among two or more datasets, and generating bootstrap network plots. The PyBootNet software package performs bootstrapping analysis on one or more feature count tables and calculates the mean and confidence intervals for seven different network metrics for every dataset. The package also provides statistical analyses such as box and whisker plots and a binomial test to determine whether individual network metrics between two different datasets are statistically different from one another. Finally, PyBootNet produces network visualizations indicating the highly supported edges and outputs the nodes with the greatest connectivity. We demonstrate the use of PyBootNet and its features with two different microbial community datasets: one from a mouse microbiome study and the other from a microbiome study of the built environment. While both examples used here to test the functionality of PyBootNet come from microbial communities, network analysis has also been applied to investigate many different biological systems at different level of complexity from entire communities to individual cells, using many types of datasets including transcription profiles, gene function abundances, and metabolomics. Thus, PyBootNet is an intuitive and easy-to-install Python package that will enable researchers to test hypotheses of network robustness and architecture for any given correlation based biological data.

## MATERIALS AND METHODS

### Data Preparation

Two sets of data were used for this project to showcase the capabilities and performance of the software. The first data set was a mouse gut microbiome data [17], while the second was a built environment bacterial dataset available at https://github.com/kaelyn20/Kelley-Lab-Projects. The software *PyBootNet* and instructions can be found on GitHub: https://github.com/Shayan-Akhavan/PyBootNet.git. The software was developed to take input data into Python 3.10.11 functions designed to operate with Pandas DataFrames. The features for network visualization are represented as separate columns in the DataFrame. The *map_columns* function improves the readability of column values by transforming their names to a standardized format of ’X1’, ’X2’, ’X3’, etc. The function maintains a dictionary that maps these transformed column names to their corresponding taxonomic information, enabling the retrieval of species-level taxonomy for each column. Next, the data was preprocessed by removing any unnecessary columns that contain numerical values. For the built environment data, “ATP”, “WetMat”, “Sample”, “Objects/mL”, “Objects/cm2”, “gDNA well”, “gDNA plate”, “spore_density”, “hyphae_length” were removed due to their values are unnecessary for correlation calculation. For the mouse data, “Mouse_Label_Cols”, “Mouse”, “Weight”, and “Mouse_Week” were removed. Correlation coefficients were calculated using centered log-ratio (clr) transformed abundances for both the mouse and built environment datasets to account for the inherent compositionality of next-generation sequencing data.

**Table 1.** List of functions from *PyBootNet* to perform analysis.

| Functions | Description |
| --- | --- |
| map_columns() | Changes the values in a specified column of a DataFrame to be more readable, mapping them to 'X1', 'X2', 'X3', etc. |
| bootstrap_replicates() | Creates bootstrap replicates of the input data. The default number of iterations is 100. |
| correlation_matrix() | Calculates the Spearman correlation matrix for each DataFrame in the input list, taking in only numerical values. |
| calculate_network_statistics() | Calculates network statistics for each correlation matrix in the input list, using a specified threshold. |
| analyze_network_statistics() | Analyzes the network statistics for different projects, calculates descriptive statistics, and creates box plots for each statistic. |
| build_network_graph() | Builds a network graph from the input correlation matrices, averaging them if multiple matrices are provided. |
| top_nodes() | Identifies the top nodes with the highest degree in the network graph. |
| net_stat_binomial_test() | Performs a binomial test on the network statistics of two sets of bootstrap replicates. Takes the outputs of the calculate_network_statistics. |

### Bootstrapping

The *bootstrap_replicates* function generates bootstrap replicates of the data, with a default parameter of 100 iterations. In each iteration, the function randomly selects samples from the feature table with replacement from the original data (Figure 1). In the case of microbial communities, a “sample” consists of a numerical vector with each element of the vector is a count estimate of a given bacteria in that sample. For example, a mouse gut sample would comprise counts of all the bacterial genera identified via next generation sequencing for a single fecal sample. For the mouse gut microbiome dataset which had a total or 74 samples, we generated 5000 bootstrap replicates for each network analysis. For the larger built environment which had a total of 238 samples, we generated 500 bootstrap replicates for each network analysis. The resulting bootstrap replicates were stored as matrices in a list, which were used to calculate the correlations within each matrix.

**Figure 1.**
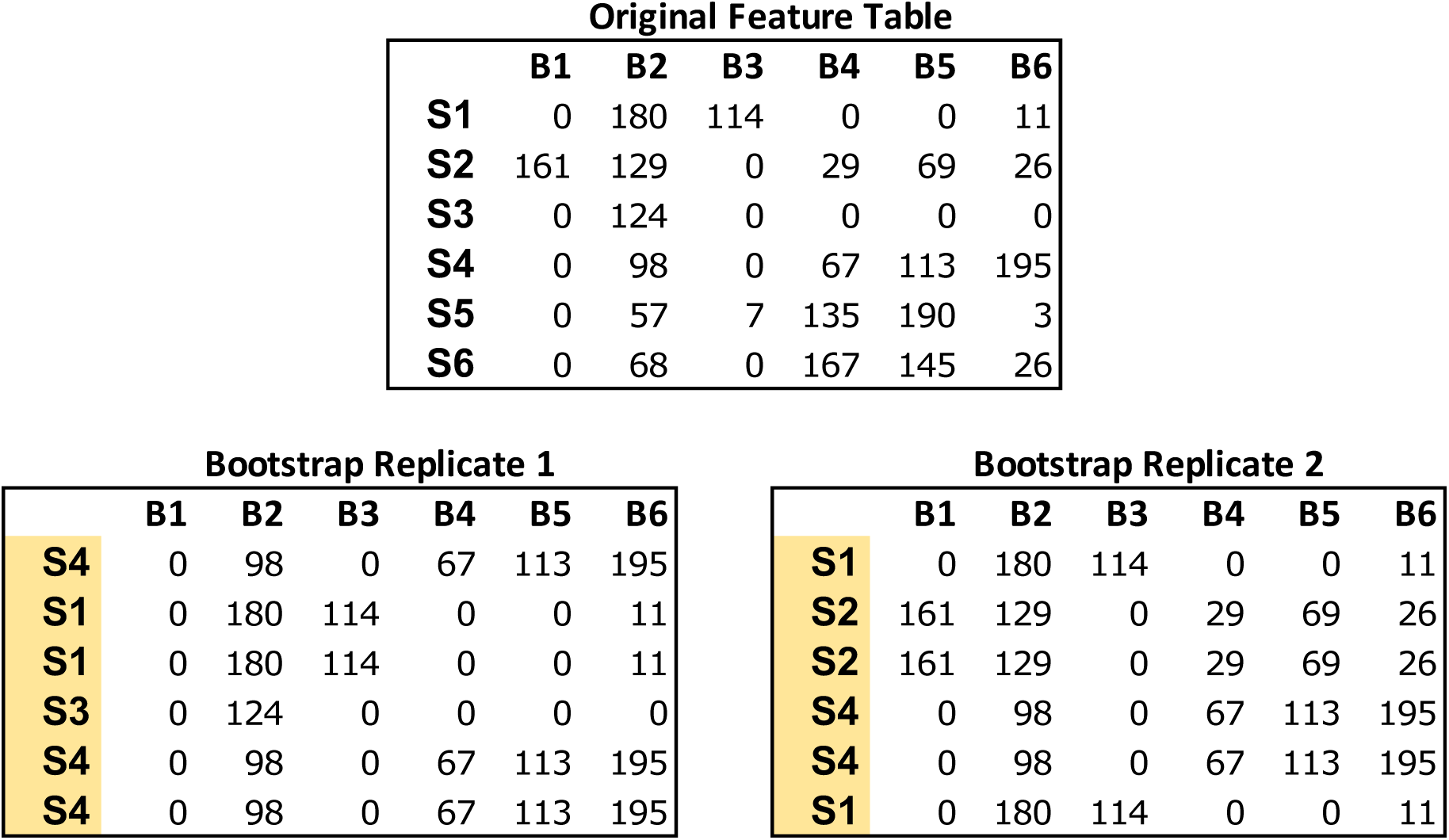
Example of bootstrapping an input feature table data of microbial count data. The Original Feature Table at the top includes 6 samples (S1-S6) and counts of 6 bacterial taxa (B1-B6). The two bootstrap replicates at the bottom are the same size as the original data (6X6) but the samples have been resampled with replacement. Any numerical feature table can be used with PyBootNet. For the test data we used the clr-transformed versions of the feature tables.

### Correlation Matrix and Network Statistics

Spearman correlation matrices were calculated for each bootstrap to replicate using the *correlation_matrix* function, considering only numerical values after removing the columns with metadata. The *calculate_network_statistics* function was used for each correlation matrix to compute various network statistics, using a specified correlation threshold where the default parameter is 0.8 or higher for the strong correlations. This included the negative correlations that are -0.8 or lower. If the user only wants to indicate the positive correlations, the supplemental analysis has a positive function. Otherwise, the desired correlations are used to construct networks to calculate the following network statistics: number of edges, number of nodes, average degree centrality, transitivity, closeness centrality, betweenness centrality, and density. Each metric was stored in a dictionary for each replicate sample and dictionary was appended to a list where the final output is a list of all the bootstraps.

### Network Analysis and Visualization

The *analyze_network_statistics* function compares network statistics across different projects. It takes a list of corresponding dictionaries of network statistics as input and generates comprehensive statistics of the data. The function calculates descriptive statistics, the mean and standard error using standard deviation, for each network statistic within each project. These results are stored in a Python pandas data frame, which is then saved as a CSV file for easy access and further inspection. Additionally, the function creates box and whisker plots for each network statistic, allowing for visual comparison of the values across different projects. These plots are automatically saved as SVG extension image files and displayed for immediate interpretation. By providing a concise summary table and intuitive visualizations, the *analyze_network_statistics* function facilitates the exploration and understanding of network statistics across multiple projects, enabling researchers to draw meaningful conclusions and make data-driven decisions.

The *build_network_graph* function constructs a network graph based on correlation matrices, providing a visual representation of the relationships between features. It accepts either a single correlation matrix or a list of correlation matrices as input. If a list is provided, the function averages the matrices to obtain a single consolidated matrix. The function then creates a network graph using the NetworkX library [18], where each variable is represented as a node, and the correlations between variables are represented as weighted edges. The magnitude and sign of the correlation values determine the strength and direction of the correlations. The function allows for the filtering of significant correlations based on a specified threshold value. The resulting network graph is visualized using Matplotlib [18], with nodes colored in sky blue and edges colored in red for negative correlations and blue for positive correlations. The edge thickness is proportional to the absolute value of the correlation. A legend is created to differentiate between positive and negative correlations. The graph is saved as an SVG file and displayed for visual inspection. This function provides researchers with a powerful tool to explore and understand the complex relationships between variables in their data, facilitating the identification of key connections and potential patterns. The *top_nodes* function identified the nodes with the highest degree in the network graph, while the *most_connected_nodes* function determined the most connected node. The *nodes_edges_table* function created a Table of the number of edges for each node.

The PyBootNet workflow for each bootstrap replicate is illustrated in Figure 2.

**Figure 2.**
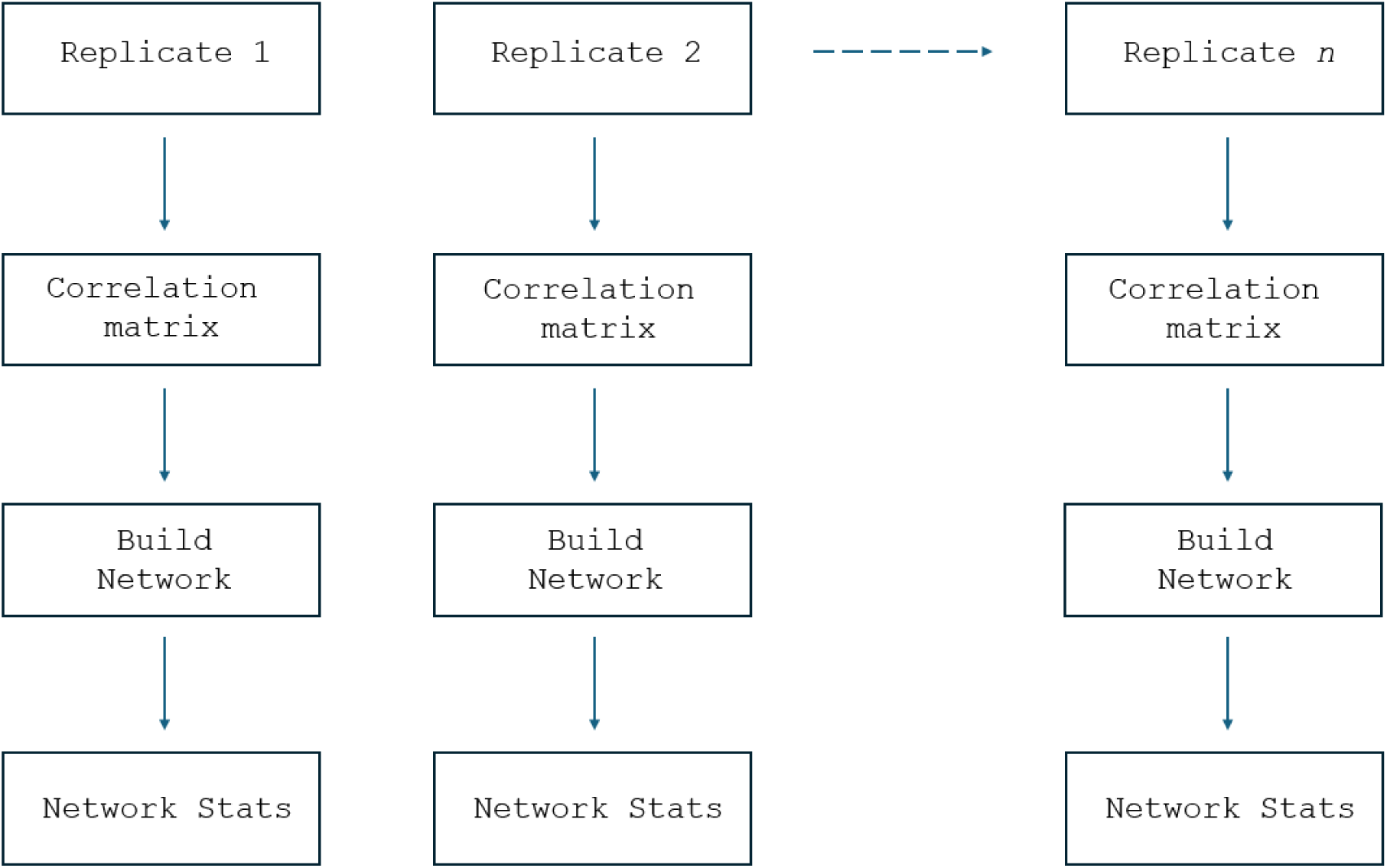
PyBootNet workflow of creating replicates and calculating individual correlations by their respective newly formed datasets. Network constructions follow and stats are calculated.

### Data Export and Statistical Testing

The *save_table_to_csv* function was implemented to store preprocessed data as a CSV file for import. A binomial test was performed on the network statistics of two sets of bootstrap replicates using the *net_stat_binomial_test* function, which takes the outputs of the *calculate_network_statistics* function. The binomial test is conducted using a list, where each element of the list is a dictionary containing the calculated network statistics.

### Additional Analyses

The *build_positive_network* function constructs a network graph focusing on positive correlations above a specified threshold, while the *build_filtered_networks* function builds a filtered network graph, retaining only nodes with a maximum degree specified by the parameter *max_degree*. The *build_negative_networks* function creates a network graph focusing on negative correlations below a specified threshold, and the *negative_filtered_networks* function builds a filtered network graph of negative correlations, where both parameters have the same values. The b*ootstrap_sample_with_correlation* function performs bootstrapping on the input data, calculates correlations for each bootstrap replicate, and returns the average correlation matrix. Finally, the *top_nodes_network_graph* function constructed a network graph highlighting the top nodes with the highest degree of centrality. The default parameter is set to 20 nodes, but the user can adjust accordingly. All functions were designed to provide flexibility in input data format and output file formats, facilitating integration into the analysis pipeline. Input parameters and file names were adjusted according to the specific use case of the thesis.

### Computational Testing Platform

*PyBootNet* analysis was run on Windows 10 Home 64-bit operating system with an AMD Ryzen 7 5800X 8-core processor at 3.8 GHz clock speed. The dedicated memory available was 32 gigabytes and the graphical processor was an Nvidia GeForce RTX 3080 with 10 gigabytes of video memory.

## RESULTS AND DISCUSSION

### Mouse Data

Eight sub-datasets were constructed from mouse microbiome data (Sisk-Hackworth et al., https://doi.org/10.1101/2024.07.01.601610) from fecal bacterial libraries collected from 4 different groups at two different time points, Week 3 and Week 10. The four groups included two different genotypes, wild type and mutant, from both sexes, female and male, and were based on 20 samples per group. Spearman rank correlation [20] values were calculated for each sub-dataset using 5000 bootstrap replicates, with a computation time of about 2 minutes per dataset. For each set of correlations, network statistics were calculated. The computed network properties for each bootstrap replicate with mean r correlations values of 0.8 or greater included the number of edges, number of nodes, average degree centrality, transitivity, closeness centrality, betweenness centrality, and density. The calculation of these network features for each 177x177 pairwise correlation matrix required approximately 10 minutes and 20 seconds of computational time per feature. After collecting and storing the network statistics, the process of visualizing the networks for each mouse type was completed in approximately 10 seconds. The software is designed to be user-friendly and efficient in analyzing the mouse gut microbiome data. After ensuring that the input data fit the required input format for the functions, the analysis can be performed by simply executing the relevant functions provided by the software.

The *analyze_network_statistics* function was executed to calculate and compare the network properties of the gut microbiome data from Week 10 male wild type and male mutant type mice. The function took 1.0 second to run, with the majority of the computational time being spent on the *calculate_network_statistics_ function*. This function computes the mean and standard error of the mean for each network statistic across all bootstrap replicates. The software is designed to handle multiple conditions, appending additional columns to the output for each condition, facilitating the comparison of network statistics across different groups (Figure 3).

**Figure 3.**
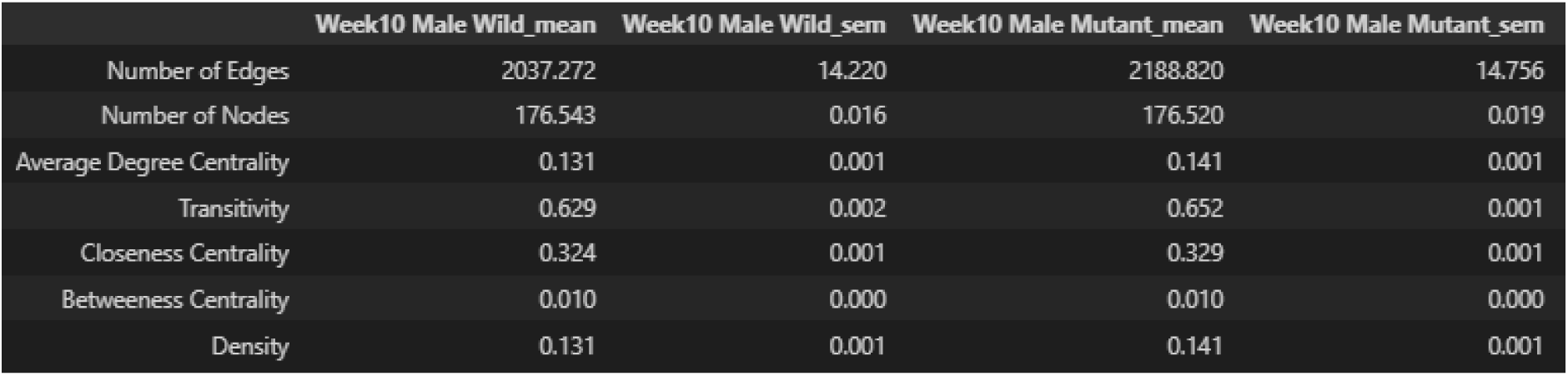
Example raw output showing network statistics for fecal bacterial communities collected from two different genotype groups of male mice (n=20/group), wild type and mutant, at week 10 using the *analyze_network_statistics* function. For each group, means and standard error (sem) were calculated for 5 network metrics.

The *analyze_network_statistics* function, which generates the output shown in Figure 4, also produces associated box and whiskers plots that are saved as .PNG image files by default. Figure 4 showcases two plots that highlight distinct differences between the female wild type and other mouse types in terms of transitivity and betweenness centrality. The female wild type exhibits lower transitivity and higher betweenness centrality compared to the other groups. These findings suggest that the clusters within the female wild type of gut microbiome network are more interconnected and have a greater influence on the overall network structure[22].

**Figure 4.**
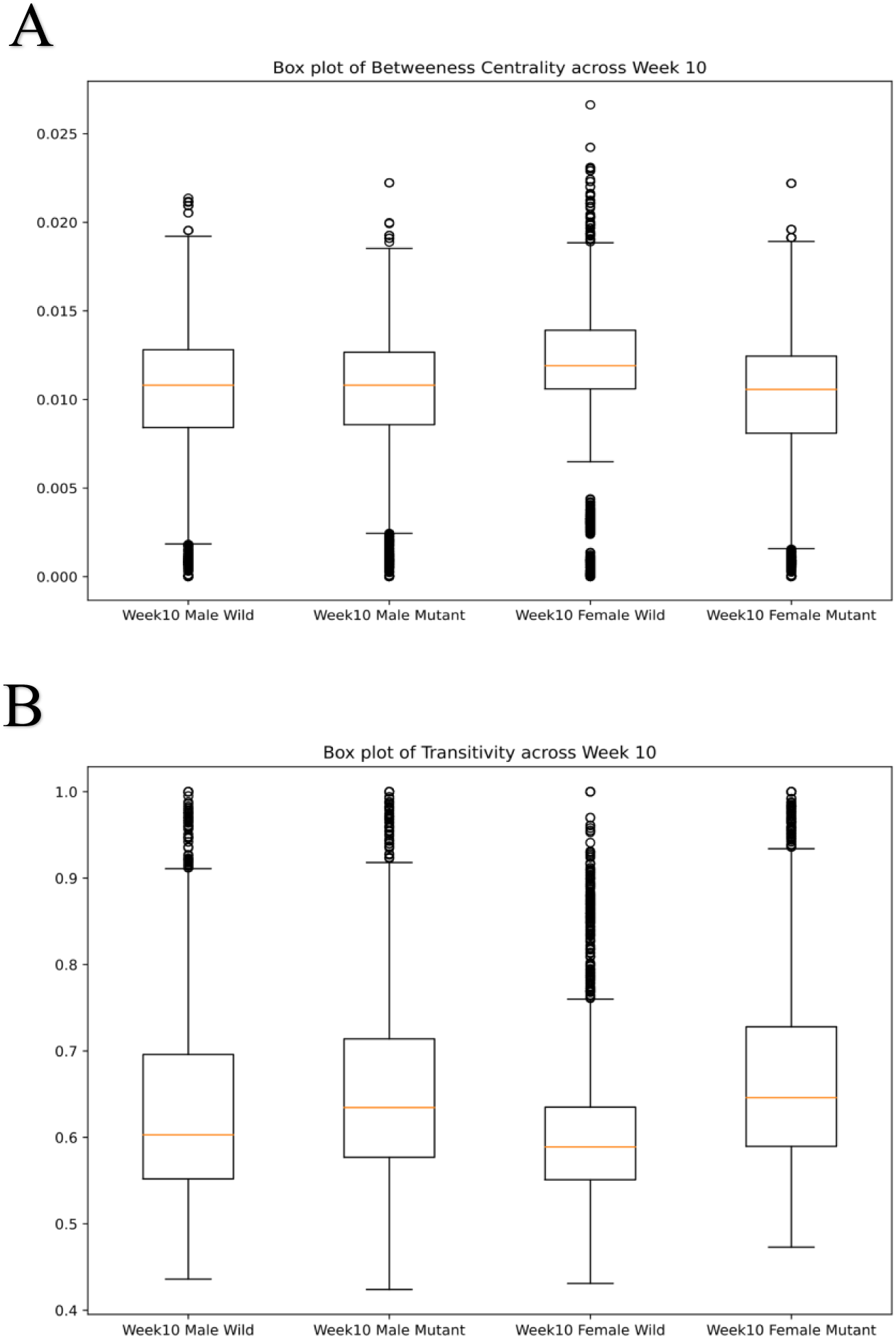
Box and Whiskers plot comparing betweenness centrality (A) and transitivity (B) among networks of the 4 mouse groups at Week 10.

The software also includes a binomial test function that compares the network statistics of two projects, as illustrated in Figure 5. This functionality enables researchers to conduct comparative analyses between different conditions or time points, such as the one presented in Table 2. The table showcases the results of binomial bootstrap tests (n=5000 replicates) comparing the network statistics of gut microbiome data from male wild type mice at Week 3 and Week 10, as well as male wild type and male mutant type mice at Week 10. The output provides valuable insights into the differences in network properties between the compared groups, aiding in the interpretation of the complex interactions within the mouse gut microbiome.

**Figure 5.**
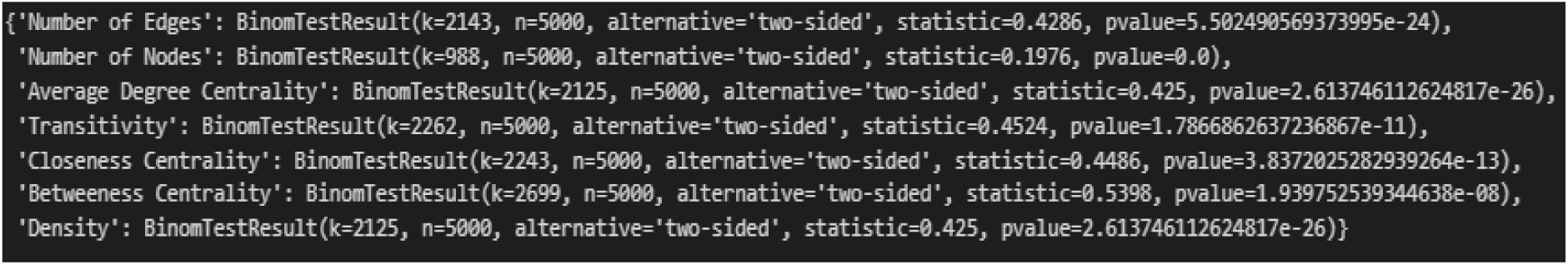
Example raw output generated by *net_stat_binomial_test()* comparing male wild type Week 3 and Week 10 network metrics.

**Table 2.** Binomial bootstrap tests results (n=5000 replicates) comparing Week 3 to Week 10 male wild type networks, and male wild type to male mutant type networks at Week 10.

| Male Wild Type | Week 3 | Week 10 | k | statistic | p-value |
| --- | --- | --- | --- | --- | --- |
| Number of Edges | 186814.0 | 2037.3±14.2 | 2143 | 0.4286 | 5.50e-24 |
| Number of Nodes | 176.60.02 | 176.5±0.02 | 988 | 0.1976 | 0.00* |
| Average Degree Centrality | 0.120.001 | 0.13±0.001 | 2125 | 0.425 | 2.61e-26 |
| Transitivity | 0.6140.001 | 0.629±0.002 | 2262 | 0.4524 | 1.79e-11 |
| Closeness Centrality | 0.3170.001 | 0.324±0.001 | 2243 | 0.4486 | 3.84e-13 |
| Betweenness Centrality | 0.010.0 | 0.01±0.0 | 2699 | 0.5398 | 1.94e-08 |
| Density | 0.120.001 | 0.131±0.001 | 2125 | 0.425 | 2.61e-26 |
| Week 10 | Male Wild | Male Mutant | k | statistic | p-value |
| Number of Edges | 2037.3±14.2 | 2188.8±14.8 | 2442 | 0.4884 | 0.104 |
| Number of Nodes | 176.5±0.02 | 176.5±0.02 | 924 | 0.1848 | 0.0* |
| Average Degree Centrality | 0.13±0.001 | 0.141±0.001 | 2212 | 0.4424 | 3.93e-16 |
| Transitivity | 0.629±0.002 | 0.652±0.001 | 2098 | 0.4196 | 5.23e-30 |
| Closeness Centrality | 0.324±0.001 | 0.329±0.001 | 2380 | 0.476 | 7.23e-4 |
| Betweenness Centrality | 0.01±0.0 | 0.01±0.0 | 2592 | 0.5184 | 0.009 |
| Density | 0.131±0.001 | 0.141±0.001 | 2212 | 0.4424 | 3.93e-16cc |

\* p-value less than 1.0e-30.

Figure 6 shows PyBootNet visualizations of bootstrapped networks based on samples from two different groups of mice with nodes, edges, and labels. Identifying the bacterial communities with the highest number of nodes can provide valuable insights into the key players within the gut microbiome under different conditions. By analyzing the top nodes, researchers can formulate hypotheses about the interactions and dynamics occurring between these specific bacterial groups. Figure 7 showcases the top 3 taxa from each week 10 mouse type, as determined by the *top_nodes* function. This information can serve as a starting point for further investigations into the roles and relationships of these prominent bacterial communities within the gut microbiome. Furthermore, the analysis reveals that the top bacterial taxa vary between different mouse types, suggesting that the growth and composition of the gut microbiome may be influenced by sex-linked factors and the presence or absence of specific mutations. These findings potentially highlight the importance of considering both genetic background and sex when investigating the complex interactions within the gut microbiome.

**Figure 6.**
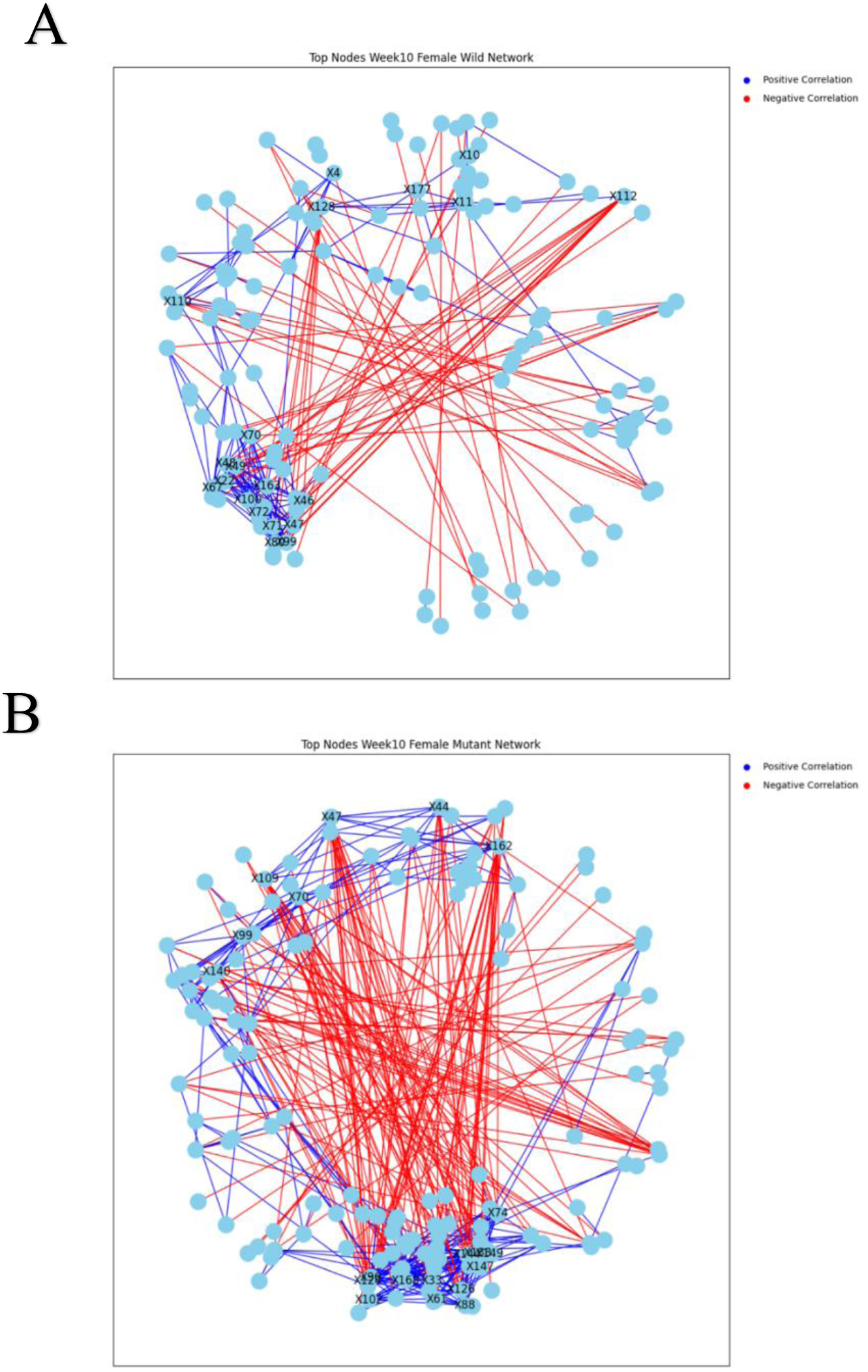
Mutant (A) and Female Wild Type (B) network visualizations indicating positive (blue) and negative (red) lines correlations. The top 20 nodes with highest degree are labels on the network (e.g., X10) using the *top_nodes_network_graph()* function.

**Figure 7.**
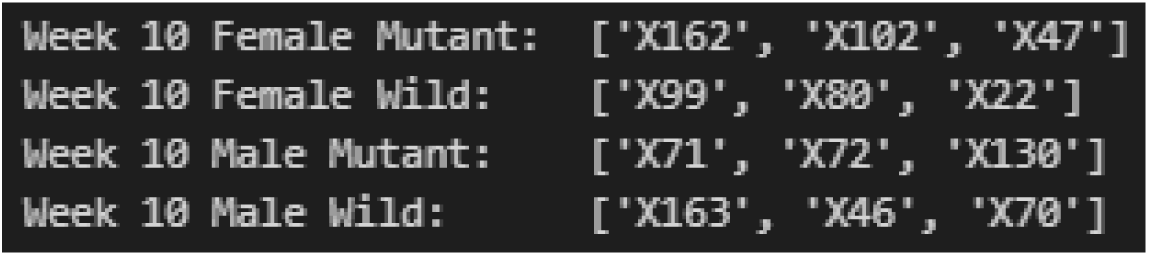
Example of raw output of the use of the *top_nodes* function, with the parameter set to show 3 of the nodes with highest degrees.

### Built Environment Data

Six sub-datasets were constructed from the built environment data [17] by their material type and treatment condition. Generating correlation matrices from the bootstrap replicates was considerably faster for the mouse data than for the built environment data, despite the latter having a larger 361x361 matrix compared to the 177x177 matrix of the mouse data. The generation of the bootstrap replicates did not different markedly between the datasets. However, computing the network statistics for each correlation matrix for the build environment dataset took approximately 10 minutes. The increased processing time observed when analyzing the built environment data can be attributed to the larger matrix size, which requires more memory caching during the computation of each network statistic. Despite this increase in computational requirements, the same functions used for the mouse microbiome data were successfully applied to the built environment dataset, allowing for the identification of differences between the various conditions.

The descriptive statistics of the network metrics facilitate comparisons between the wet and cycling treatment conditions, as well as between different building materials subjected to a specific treatment condition. Figure 8 shows the PyBootNet output of the descriptive network statistics for samples from three built material datasets. The pairwise binomial tests found no significant differences between wet gypsum and cycling gypsum (within material; raw output Figure 9) but highly significant differences between different materials under the same conditions (Table 3). There were also clear differences between wet and cycling when these were combined from all material types (Figure 10). PyBootNet’s ability to efficiently handle larger datasets of hundreds of samples on a laptop computer enables researchers to perform these calculations without encountering computational difficulties, facilitating the analysis of complex microbial communities in various environments.

**Figure 8.**
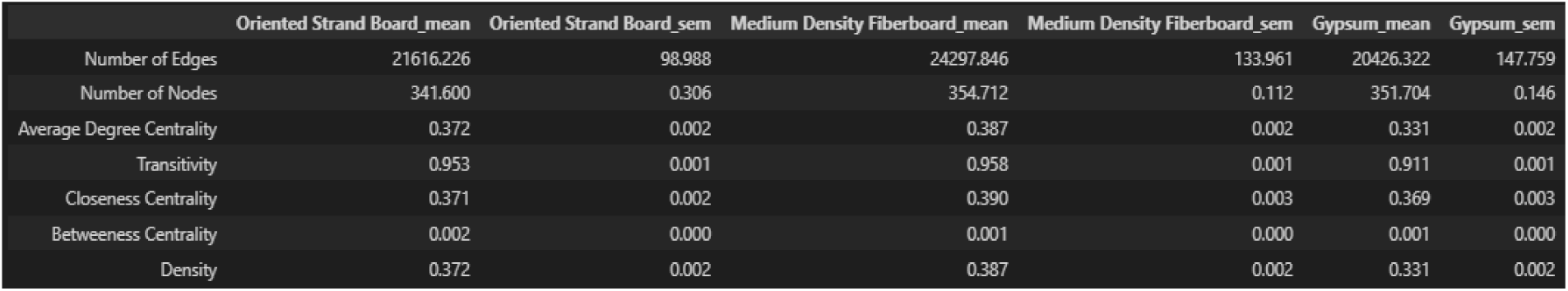
Example of raw output of network statistics (mean and standard error) for three different material types using the *analyze_network_statistics* function.

**Figure 9.**
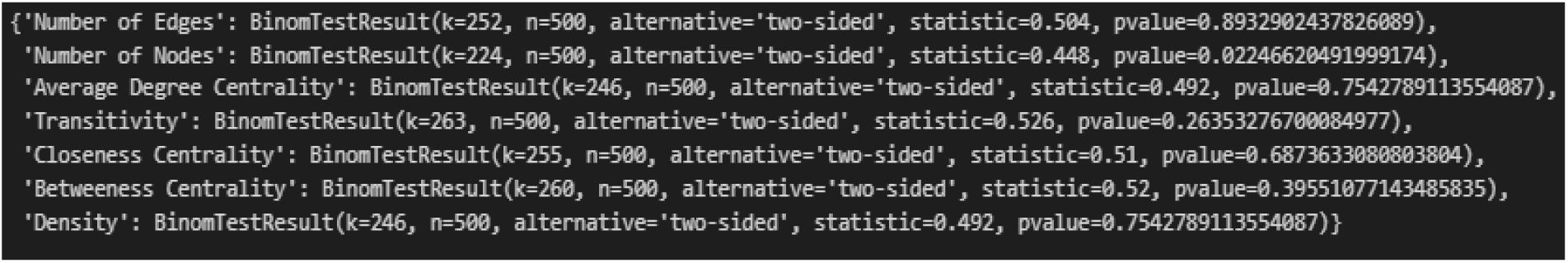
The raw output of utilizing the *net_stat_binomial_test()* for comparing gypsum materials under wet and cycling conditions

**Figure 10.**
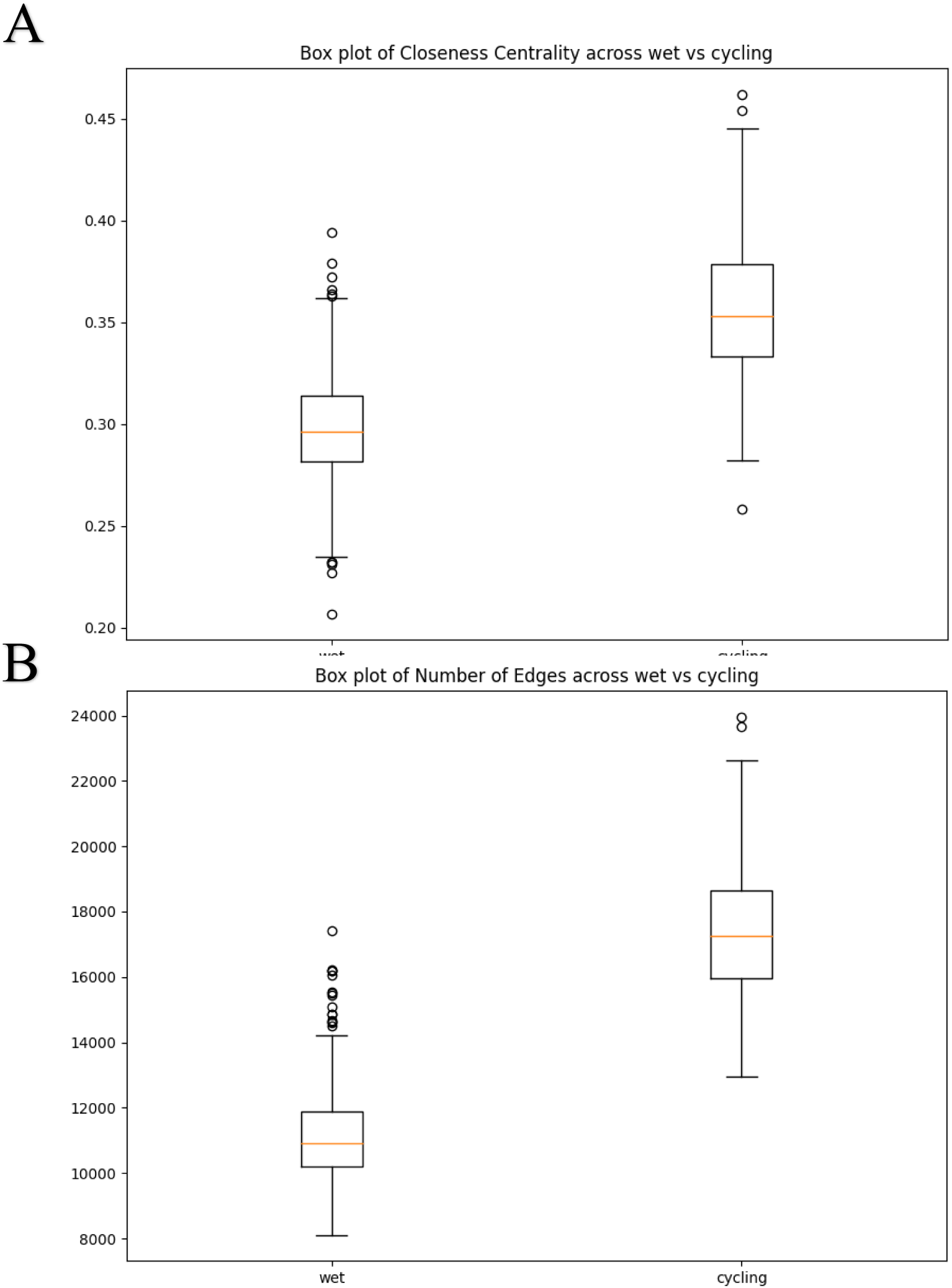
Box and Whiskers plot of number of edges (A) and closeness centrality (B) between networks based combined wet vs cycling data.

**Table 3.** Binomial bootstrap tests (n=500 replicates) comparing gypsum material under wet and cycling conditions and comparing medium density fiberboard (MDF) material and gypsum under wet conditions.

| Gypsum | Wet | Cycling | k | statistic | p-value |
| --- | --- | --- | --- | --- | --- |
| Number of Edges | 20344.4±149.4 | 20531.6±145.0 | 247 | 0.494 | 0.823 |
| Number of Nodes | 351.7±0.14 | 351.79±0.14 | 221 | 0.442 | 0.348 |
| Average Degree Centrality | 0.330±0.002 | 0.333±0.002 | 246 | 0.492 | 0.687 |
| Transitivity | 0.912±0.001 | 0.911±0.001 | 243 | 0.486 | 0.823 |
| Closeness Centrality | 0.364±0.003 | 0.368±0.003 | 247 | 0.494 | 0.964 |
| Betweenness Centrality | 0.001±0.0 | 0.001±0.0 | 181 | 0.362 | 0.561 |
| Density | 0.33±0.002 | 0.333±0.002 | 246 | 0.492 | 0.687 |

| Wet | MDF | Gypsum | k | statistic | p-value |
| --- | --- | --- | --- | --- | --- |
| Number of Edges | 24486.4±129.4 | 20344.4±149.4 | 93 | 0.186 | 7.41e-48 |
| Number of Nodes | 354.6±0.11 | 351.7±0.14 | 104 | 0.208 | 3.86e-41 |
| Average Degree Centrality | 0.391±0.002 | 0.330±0.002 | 102 | 0.204 | 2.59e-42 |
| Transitivity | 0.955±0.001 | 0.912±0.001 | 57 | 0.114 | 4.34e-75 |
| Closeness Centrality | 0.396±0.003 | 0.364±0.003 | 195 | 0.39 | 9.91e-7 |
| Betweenness Centrality | 0.001±0.0 | 0.001±0.0 | 293 | 0.586 | 1.4e-4 |
| Density | 0.391±0.002 | 0.33±0.002 | 102 | 0.204 | 2.59e-42 |

## CONCLUSION

PyBootNet provides a much needed, user-friendly and efficient Python package for bootstrapping analysis and network construction. The intuitive application of PyBootNet to two diverse microbial community datasets, a mouse gut microbiome study, and a microbiome study of the built environment, demonstrates the package’s versatility and potential for widespread use. The software’s ability to generate bootstrapped network metrics, statistically compare network metrics among datasets, and visualize bootstrap networks enables researchers to formulate data-driven hypotheses and uncover meaningful patterns in microbial ecosystems, but it can also be used to generate and statistically compare correlation networks for any type of data.

PyBootNet’s performance evaluation highlights its computational efficiency and scalability, making it suitable for analyzing datasets with hundreds of samples. The package’s minimal dependencies and straightforward input requirements further contribute to its accessibility and ease of use. The development of PyBootNet addresses the limitations of existing packages, such as BioNetComp and BootNet, by offering a more flexible and generalizable Python-based solution for bootstrapping analysis and network construction. By integrating these functionalities into a single, user-friendly package, PyBootNet streamlines the analysis process and facilitates the exploration of microbial community interactions. As the package continues to be refined and expanded, it has the potential to become an essential tool for network analysis of many biological systems

## CODE AVAILABILITY

All the code, installation instructions and test files can be found at: https://github.com/Shayan-Akhavan/pybootnet/tree/main

## ACKNOWLEDEGMENTS

We thank L. Sisk-Hackworth and K. Naninni for providing the data used to test and enhance PyBootNet’s capabilities. L. Miller provided essential guidance for calculating standard errors, C. Zúñiga guided network construction, and both provided helpful suggestions on the text. Members of the Kelley Lab offered valuable insights on developing new software and helping the lead author stay on track.

